# Reversible Actin modifications by Mical and SelR regulate dynamic actomyosin ring functions during cell wound repair

**DOI:** 10.64898/2026.07.05.736623

**Authors:** Mitsutoshi Nakamura, Justin Hui, Susan M. Parkhurst

## Abstract

Cell wound repair requires rapid and coordinated remodeling of the actin cytoskeleton to restore cortex integrity. Here, we show that a Rab35-Mical-SelR pathway regulates actomyosin ring dynamics through reversible actin redox. We find that Rab35 is recruited to wounds and is essential for proper actin ring assembly and disassembly. Rab35 regulates the recruitment of Mical, an actin-oxidizing enzyme, and SelR, a reductase that reverses oxidation, to the cell wound. Mical and SelR knockdowns disrupt actin ring formation and wound closure, whereas double knockdown partially rescues these defects, indicating a balanced redox cycle is required. Super-resolution microscopy reveals that Mical and SelR differentially regulate F-actin architecture and orientation. Mutation of actin at Methionine 44 does not fully recapitulate Mical knockdown phenotypes, suggesting the presence of additional targets and enzymes. Taken together, our results indicate that reversible actin modifications dynamically regulate F-actin architecture and orientation for actin ring assembly and disassembly.

## INTRODUCTION

Throughout normal development, the cell cortex must be continuously remodeled to form the specialized domains required to carry out essential dynamic cellular processes with high fidelity, including signal transduction/communications, cell division, migration, intracellular transport, endocytosis/exocytosis, and inter- and intra-cellular attachments/adhesion (*1–5*). This plasticity is also essential when things go awry: cells are subjected to daily assaults from their external environment that lead to cell cortex damage and/or breaches (*6–9*). When cell cortex integrity is disrupted through trauma, infection, or disease, the cell must not only rapidly close the breach, but also remodel the region involved following wound closure to remove components of the repair process (i.e., membrane plug and actomyosin ring remnants) and re-establish the original cell cortex (i.e., lipid/protein composition, organization, and attachments)—or remain forever impaired (*7–13*).

One protein known to affect aspects of cell cortex remodeling is Rab35. Rab35 belongs to the large highly conserved family of Rab GTPases (>60 members in the human genome) that are well-known regulators of membrane and vesicle trafficking (*14–18*). For example, Rab35 has been shown to mediate engulfment and clearance of dead cells by autophagy and to regulate neurite outgrowth through synaptic vesicle trafficking and turnover (*16, 19–21*). More recently, Rab family GTPases have been implicated in actin and microtubule dynamics, including the regulation of F-actin clearance at the abscission site during cytokinesis (*20, 22–26*). Like other GTPase proteins, Rab35 carries out its roles through the recruitment of different effectors that regulate non-redundant downstream pathways (*16, 20, 21*). For its roles in actin dynamics, one such Rab35 effector is Mical (Molecule interacting with CasL), a highly conserved protein encoding a multi-domain flavoprotein monooxygenase enzyme. Mical can bind directly to actin where it oxidizes two of its conserved amino acids [Methionine (Met) 44 and 47]. Oxidation of Met44 by Mical leads to actin disassembly (*27–30*). Interestingly, this actin modification is reversible. SelR (Selenoprotein R), a conserved methionine sulfoxide reductase, acts in opposition to Mical by reducing oxidized actin (*31, 32*).

Using the *Drosophila* cell wound model, we show that Rab35 and its downstream effectors, Mical and SelR, are recruited to wounds and required for efficient cell wound repair. Rab35, Mical, and SelR are all required for the assembly of a robust actomyosin ring around the wound periphery. Interestingly, both Mical and SelR knockdowns disrupt the formation of an actomyosin ring and efficient cell wound repair, but double knockdowns partially rescue the phenotypes observed in each single knockdown. Super-resolution microscopy revealed that Mical and SelR regulate F-actin architecture differently during cell wound repair, consistent with their opposite biochemical activities. Furthermore, expressing Actin M44L, a mutant actin blocking the Mical oxidation site, disrupts the formation of the actomysin ring, but does not fully mimic Mical knockdown phenotypes. Reducing Mical or SelR expression in an Actin M44L background suggests that more than one enzyme likely regulates actin oxidation, and that Mical/SelR regulates the oxidation and reduction cycle of proteins other than actin during cell wound repair. Taken together, our findings uncover reversible actin modifications by Mical and SelR as a dynamic mechanism that sculpts F-actin architecture to drive actomyosin ring assembly and disassembly at the wound periphery needed for efficient cell wound closure.

## RESULTS

### Rab35 is recruited to the wounds and is required for actin ring assembly and efficient cell wound repair

To determine what role Rab35 plays in cell wound repair, we first examined where Rab35 is recruited during cell wound repair. We generated wounds by laser ablation on the lateral side of nuclear cycle 4–6 YFP-Rab35 knock-in embryos (*33*) co-expressing an F-actin reporter (sStMCA; see Methods). In control embryos, F-actin accumulates in two distinct regions: 1) a highly enriched ring at the wound edge (actin ring) and 2) a less dense accumulation at the periphery of the actin ring (actin halo) (Fig. 1, A) (*34, 35*). Upon laser wounding, Rab35 is recruited to the actin ring and halo regions (Fig. 1, A’ and A”). While Rab35 is recruited to the actin ring region, it does not fully overlap, but rather accumulates more towards the inside of the actin ring (Fig. 1, A” and B).

**Figure 1.**
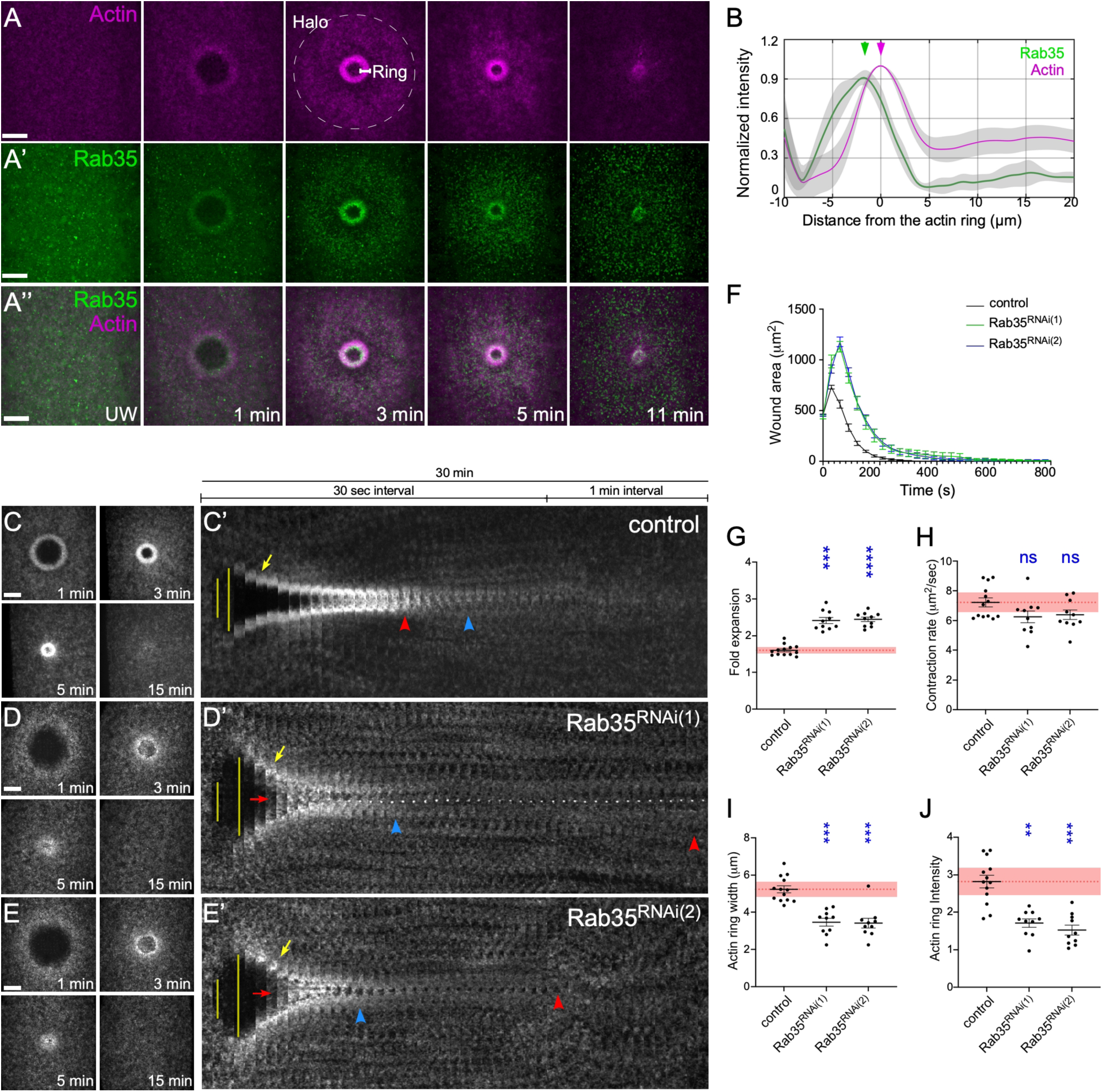
Rab35 is recruited to cell wounds and is required for actin ring dynamics and efficient wound repair. (A-A”) Confocal xy projection image from a wounded NC4–6 Drosophila embryo coexpressing YFP-Rab35 (A’-A”) and F-actin reporter (sStMCA; A, A”). (B) Normalized fluorescence intensity profile at 120 s post-wounding from 10 embryos expressing YFP-Rab35 and sStMCA. (C-E) Actin dynamics (sGMCA) during cell wound repair in NC4–6 staged Drosophila embryos: control (vermilion RNAi) (C), Rab35^RNAi(1)^ (D), and Rab35^RNAi(2)^ (E). (C’-E’) Kymographs across the wound area depicted in C–E, respectively. Wound expansion is highlighted by yellow lines. Actin recruitment to the actomyosin ring is indicated by yellow arrows. Actin accumulation internal to the wound is indicated by red arrows. Wound closure is indicated by red arrowheads. Actomyosin ring disassembly (or lack thereof) is indicated by blue arrowheads. Scale bar: 20 µm. Time after wounding is indicated. (F) Quantification of the wound area over time for control (n = 13), Rab35^RNAi(1)^ (n = 10), and Rab35^RNAi(2)^ (n = 11). (G–J) Quantification of fold wound expansion (G), contraction rate (H), actin ring width (I), and actin ring intensity (J) for control, Rab35^RNAi(1)^, and Rab35^RNAi(2)^. Error bars represent ± SEM. Kruskal-Wallis tests were performed in G–J: *P < 0.05, **P < 0.01, ***P < 0.001, ****P < 0.0001, ns is not significant.

To investigate how Rab35 affects actin dynamics during cell wound repair, we examined F-actin using an F-actin reporter (sGMCA) in embryos from two independent Rab35 RNAi knockdown backgrounds. Both RNAis efficiently knock down Rab35 (99% knockdown for RNAi(1) and 91% knockdown for RNAi(2)) and exhibit similar wound repair phenotypes. Rab35 knockdown embryos exhibit substantial F-actin remodeling defects: 1) the actin ring prematurely disassembles, 2) wounds overexpand, 3) actin ring width and intensity are decreased, and 4) prolonged F-actin accumulation is observed at the wound (Fig. 1, C-J; Table S2). These results indicate that Rab35 responds to cell wounds and functions in both actin ring assembly and disassembly during the repair process.

### Mical and SelR are recruited to cell wounds Rab35-dependently

To investigate how Rab35 regulates F-actin dynamics during cell wound repair, we focused on a downstream effector of Rab35, Mical, which regulates actin dynamics during cytokinesis (*20, 29*). Mical is a multi-domain flavoprotein monooxygenase enzyme that oxidizes actin, and is known to function in opposition to SelR, a methionine sulfoxide reductase that reduces oxidized actin (*27, 28, 31, 32*). To examine whether reduction-oxidation (Redox) reactions on actin by Mical and SelR are playing a role in actin dynamics during cell wound repair, we generated wounds in embryos co-expressing GFP-Mical or StFP-SelR with F-actin reporters (sStMCA or sGMCA, respectively). Upon laser wounding, both Mical and SelR are recruited to wounds, and their recruitment patterns overlap with the actin ring and halo (Fig. 2, A-D). Similar to Rab35 recruitment to the actin ring, both Mical and SelR accumulate more towards the inner edge of the actin ring (Fig. 2 B and D). Since Mical and SelR recruitment patterns are similar to those of Rab35, we next examined whether Mical and SelR recruitment to the wounds is regulated by Rab35. Neither Mical nor SelR are recruited to wounds in a Rab35 knockdown background (Fig. 2, E-H). Taken together, our findings indicate that Mical and SelR work downstream of Rab35 in cell wound repair.

**Figure 2.**
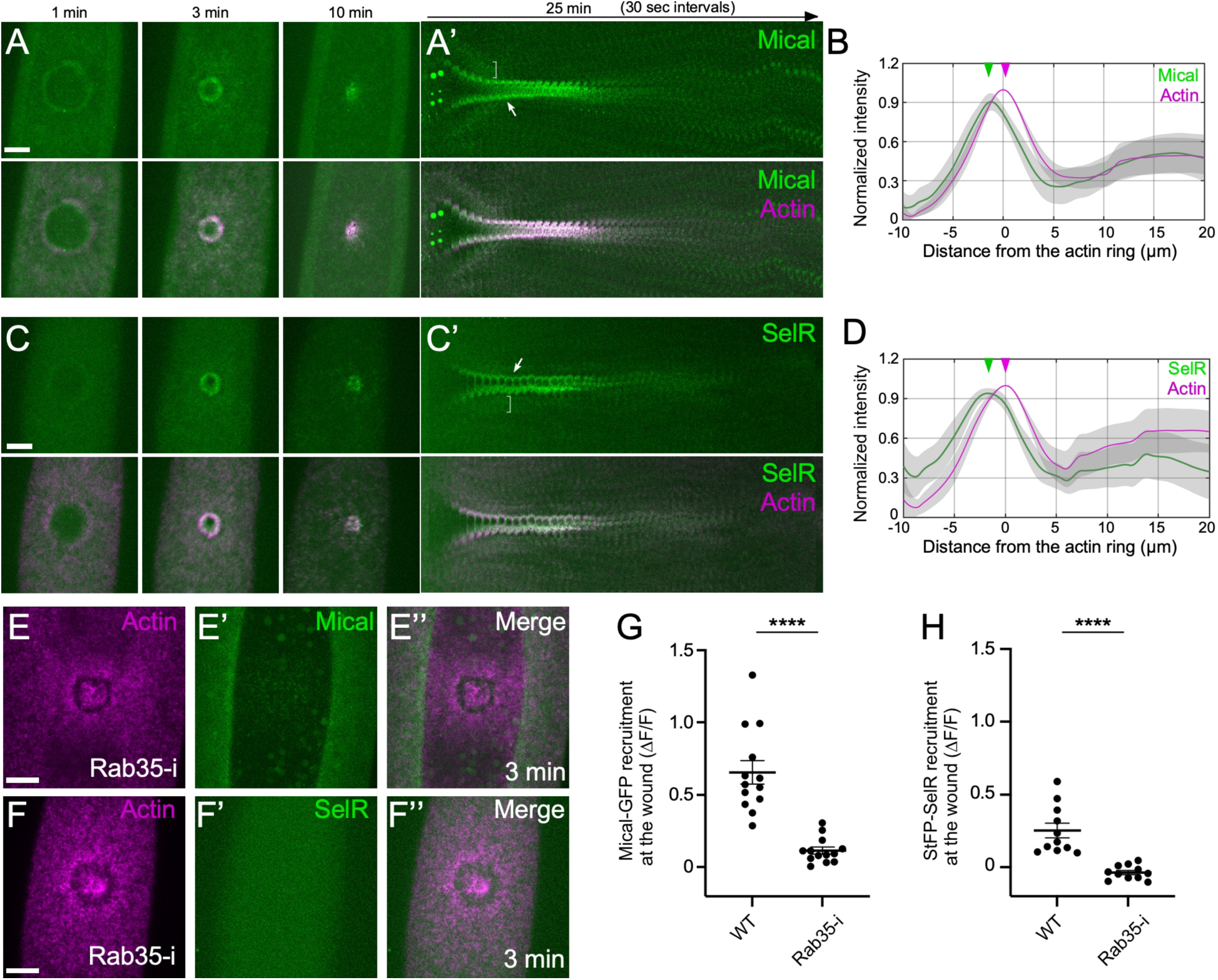
Rab35 regulates Mical and SelR recruitment to the wound. (A-A’) Confocal xy projection image and the kymograph across the wound area from a wounded NC4–6 Drosophila embryo coexpressing GFP-Mical and F-actin reporter (sStMCA). Arrow denotes ring and bracket denotes halo recruitment. (B) Normalized fluorescence intensity profile at 120 s post-wounding from 10 embryos expressing GFP-Mical and sStMCA. (C-C’) Confocal xy projection image and the kymograph across the wound area from a wounded NC4–6 Drosophila embryo coexpressing StFP-SelR and F-actin reporter (sGMCA). Arrow denotes ring and bracket denotes halo recruitment. (D) Normalized fluorescence intensity profile at 120 s post-wounding from 10 embryos expressing StFP-SelR and sGMCA. (E-F) Confocal xy projection image from a wounded NC4–6 Drosophila embryo coexpressing GFP-Mical/sStMCA (E) and StFP-SelR/sGMCA (F) in Rab35 RNAi backgrounds. (G-H) Quantification of GFP-Mical (G; n = 13 for control and n = 13 for Rab35-i backgrounds) and StFP-SelR (H; n = 11 for control and n = 11 for Rab35-i backgrounds) recruitment to the wounds at 3 min post-wounding. Scale bar: 20 µm. Time after wounding is indicated. Error bars represent ± SEM. Mann-Whitney tests were performed in G–H: ****P < 0.0001.

### Mical and SelR knockdowns disrupt actin ring assembly and disassembly during cell wound repair

To investigate the functions of Mical and SelR during cell wound repair, we generated wounds in Mical and SelR knockdown embryos expressing an F-actin reporter (sGMCA) using two independent RNAi lines with high knockdown efficiencies for each (Fig. 3A; see Methods). Intriguingly, although Mical and SelR are known to encode opposite enzymatic functions, Mical and SelR knockdowns exhibit similar wound repair phenotypes: premature actin ring disassembly, actin accumulation inside the wounds, slower wound contraction, and decreased actin ring width and intensity (Fig. 3, B-G and J-M; Table S2). To determine if Mical and SelR regulate actin dynamics differently yet result in similar phenotypes, and whether one knockdown affects the other (i.e., Mical or SelR knockdown stops the cycle of actin oxidation and reduction), we generated wounds in Mical and SelR double knockdown embryos. Surprisingly, the wound repair phenotypes in double knockdown embryos are partially rescued compared with each knockdown alone (Fig. 3H-M; Table S2), suggesting that despite their similar repair phenotypes, Mical and SelR regulate actin dynamics differently.

**Figure 3.**
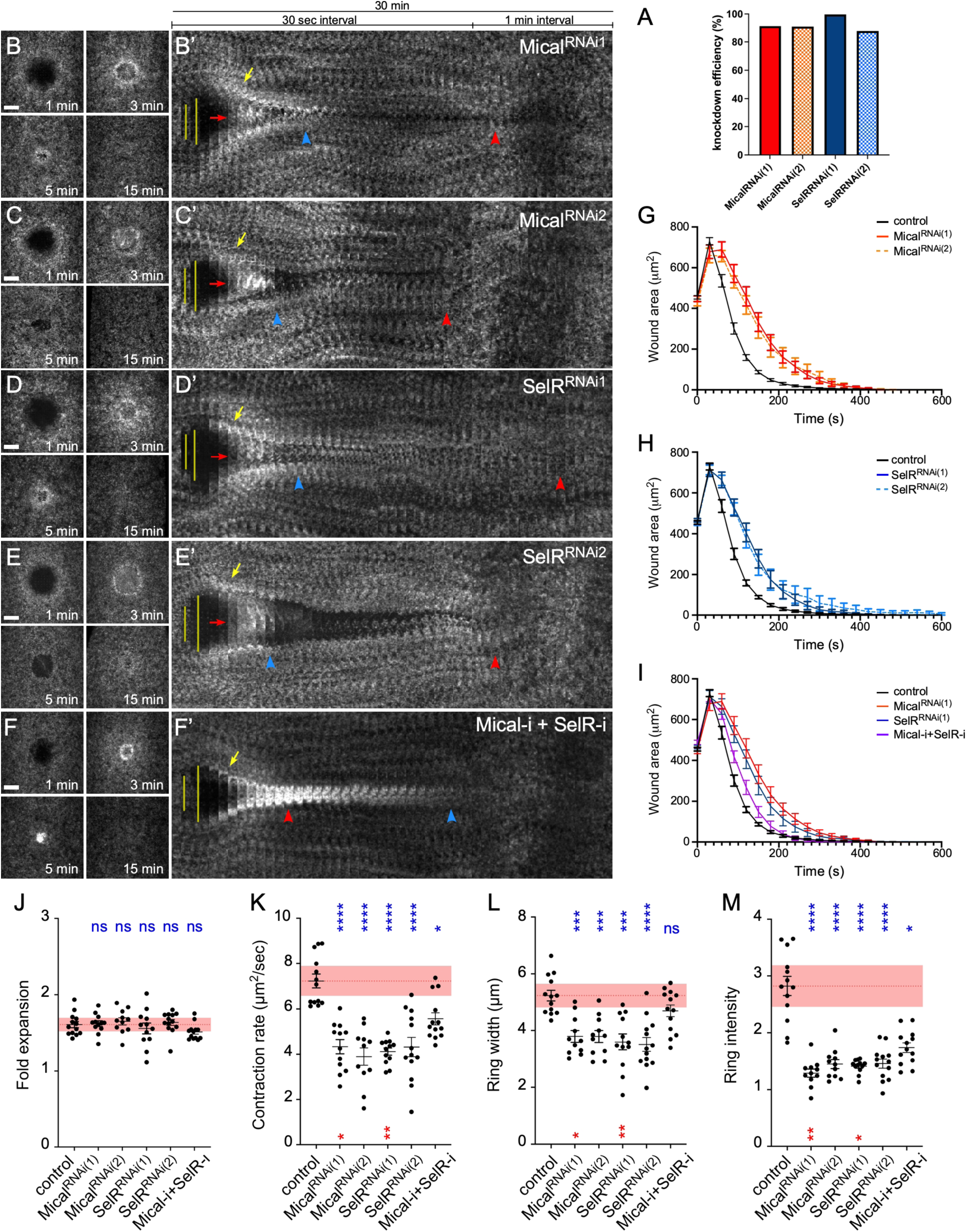
Mical and SelR are required for actin ring dynamics and efficient wound repair. (A) qPCR analysis of knockdown efficiency in Mical^RNAi(1)^, Mical^RNAi(2)^, SelR^RNAi(1)^, and SelR^RNAi(2)^ NC4–6 staged embryos. (B-F’) Actin dynamics (sGMCA) during cell wound repair in NC4–6 staged Drosophila embryos: Mical^RNAi(1)^ (B), Mical^RNAi(2)^ (C), SelR^RNAi(1)^ (D), SelR^RNAi(2)^ (E), and Mical-i+SelR-i (Mical^RNAi(1)^ and SelR^RNAi(1)^) (F). (B’-F’) Kymographs across the wound area depicted in B–F, respectively. Wound expansion is highlighted by yellow lines. Actin recruitment to the actomyosin ring is indicated by yellow arrows. Actin accumulation internal to the wound is indicated by red arrows. Wound closure is indicated by red arrowheads. Actomyosin ring disassembly (or lack thereof) is indicated by blue arrowheads. (G) Quantification of the wound area over time for control (n = 13), Mical^RNAi(1)^ (n = 11), and Mical^RNAi(2)^ (n = 11). (H) Quantification of the wound area over time for control (n = 13), SelR^RNAi(1)^ (n = 12), and SelR^RNAi(2)^ (n = 13). (I) Quantification of the wound area over time for control (n = 13), Mical^RNAi(1)^ (n = 11), and SelR^RNAi(1)^ (n = 12), and Mical-i+SelR-i (n = 13). (J–M) Quantification of fold wound expansion (J), contraction rate (K), actin ring width (L), and actin ring intensity (M) for control, Mical^RNAi(1)^, Mical^RNAi(2)^, SelR^RNAi(1)^, SelR^RNAi(2)^, and Mical-i+SelR-i. Error bars represent ± SEM. Kruskal-Wallis tests were performed in I–L: *P < 0.05, **P < 0.01, ***P < 0.001, ****P < 0.0001, and ns is not significant. Blue labels indicate comparison to control. Red labels indicate comparison to Mical-i+SelR-i. Scale bar: 20 µm. Time after wounding is indicated.

### Mical and SelR knockdowns alter F-actin architecture differently

The double knockdown phenotypes of Mical+SelR suggest that F-actin structures in the actin ring are different in each knockdown background alone. Since Mical induces F-actin disassembly through oxidizing actin and SelR allows F-actin assembly through reducing oxidized actin, we speculated that F-actin architectures in the actin ring are different between Mical and SelR knockdowns. We used super-resolution microscopy to examine F-actin architectures in Mical and SelR knockdowns at 25% and 50% wound closure (i.e., 25% and 50% of the maximum wound area) (Fig. 4; Table S3). We quantified Haralick features and F-actin orientation in the actin ring among Mical, SelR, and Mical+SelR knockdown embryos, which allows us to compare the properties of F-actin structures in these different backgrounds (*36*). Consistent with the measurements of actin ring intensity from confocal images (Fig. 3, M), the actin rings in Mical and SelR knockdown embryos are disrupted compared to control, but their F-actin architectures in the actin ring are not identical (Fig. 4, A-C and F-H). As shown earlier (Fig. 3, H-H’ and M), the formation of the actin ring in the double knockdown is more robust than in the Mical and SelR knockdowns, but less robust than in the control. (Fig. 4, A-I). To quantify differences in F-actin architecture across the different knockdown backgrounds, we performed texture analysis using Haralick features (VARIANCE, CORRELATION, UNIFORMITY, and HOMOGENEITY; see Materials and Methods) and actin orientation analysis. At 25% wound closure, the actin rings in Mical knockdown embryos exhibit a significant reduction in VARIANCE and a significant increase in CORRELATION, UNIFORMITY, and HOMOGENEITY (Fig. 4, K-N). On the other hand, the actin rings in SelR knockdown embryos exhibit a significant reduction in VARIANCE and a significant increase in UNIFORMITY and HOMOGENEITY (Fig. 4, K-N). At 50% wound closure, the actin rings in Mical and SelR knockdown embryos exhibit a significant increase in UNIFORMITY and HOMOGENEITY (Fig. 4, K-N). As Mical and SelR double knockdown partially rescues actin ring width and intensity (Fig 3 K and L), Haralick features in double knockdown embryos are partially rescued (Fig. 4, K-L). Intriguingly, while control, SelR, and Mical/SelR knockdown embryos exhibit uniform distribution of F-actin orientation in the actin ring, Mical knockdown embryos exhibit a significant increase of F-actin with Dorsal-Ventral (DV) axis orientation (Fig. 4, O-V). Taken together, these results indicate that Mical and SelR have independent functions to regulate F-actin architectures in the actin ring during cell wound repair.

**Figure 4.**
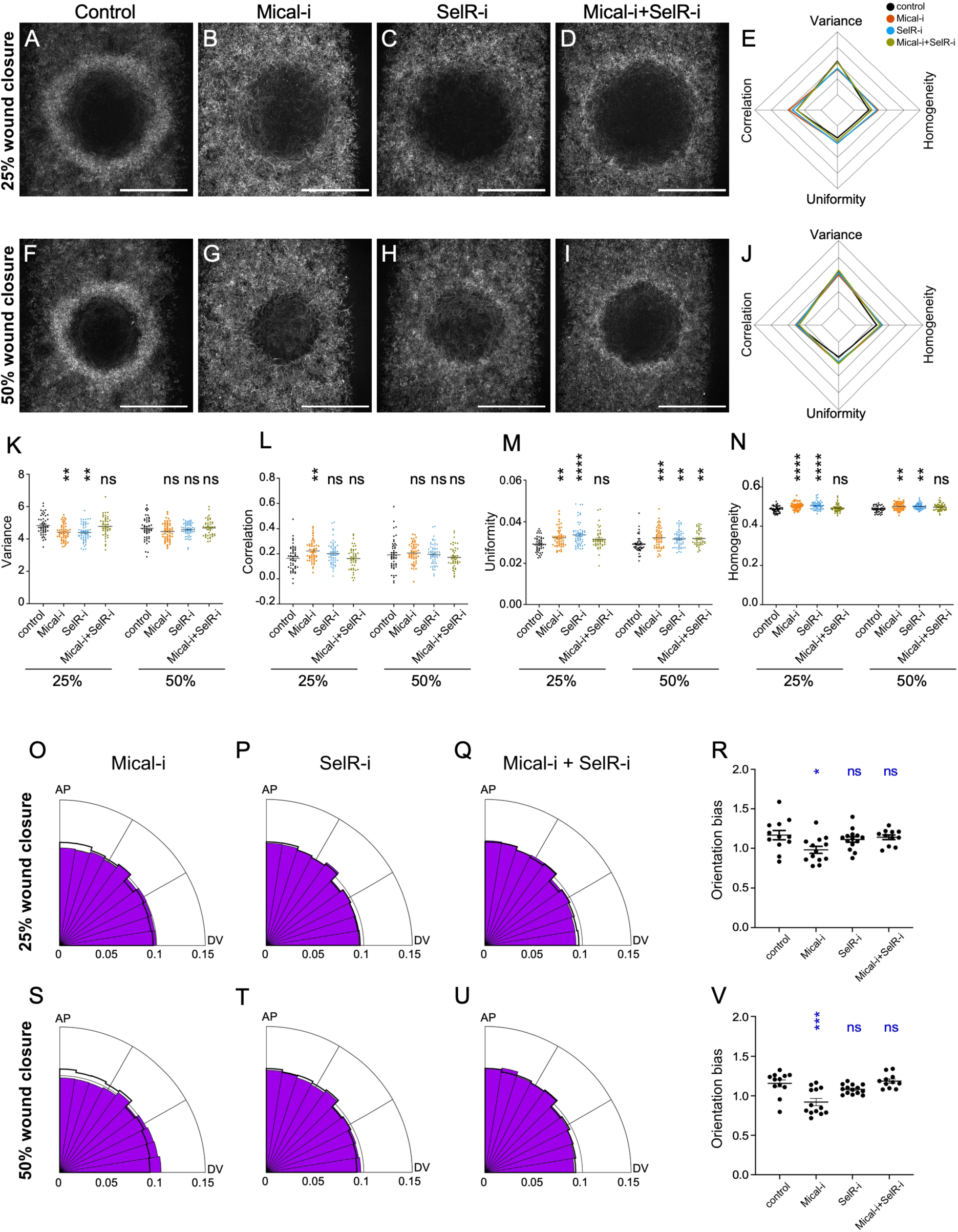
Mical and SelR knockdowns exhibit different F-acin architectures at the wound periphery. (A-D) Deconvolved super-resolution confocal images from NC4–6 Drosophila embryos expressing sGMCA at the time point corresponding to 25% of max wound area: control (A), Mical-i (B), SelR-i (C), and Mical-i + SelR-i (D). (E) Radar chart of average Haralick features comparing control, Mical-i, SelR-i, and Mical-i + SelR-i at the time point corresponding to 25% of max wound area. (F-I) Deconvolved super-resolution confocal images from NC4–6 Drosophila embryos expressing sGMCA at the time point corresponding to 50% of max wound area: control (F), Mical-i (G), SelR-i (H), and Mical-i + SelR-i (I). (J) Radar chart of average Haralick features comparing control, Mical-i, SelR-i, and Mical-i + SelR-i at the time point with 50% of max wound area. (K-N) Quantification of Haralick features comparing control, Mical-i, SelR-i, and Mical-i + SelR-i at the time points corresponding to 25% and 50% of max wound area. (O-Q) Radial histograms showing the distribution of actin filament orientations ranging from anterior–posterior (AP) bias to dorsal–ventral (DV) bias at the time point corresponding to 25% of max wound area: Mical-i (O), SelR-i (P), and Mical-i + SelR-i (Q). The black line indicates the control distribution. (R) Quantification of the distribution of actin filament orientations comparing control, Mical-i, SelR-i, and Mical-i + SelR-i at the time point corresponding to 25% of max wound area. (S-U) Radial histograms showing the distribution of actin filament orientations ranging from anterior–posterior (AP) bias to dorsal–ventral (DV) bias at the time point corresponding to 25% of max wound area: Mical-i (S), SelR-i (T), and Mical-i + SelR-i (U). The black line indicates the control distribution. (V) Quantification of the distribution of actin filament orientations comparing control, Mical-i, SelR-i, and Mical-i + SelR-i at the time point corresponding to 25% of max wound area. Scale bar: 20 µm. 12 (control), 13 (Mical-i), 14 (SelR-i), and 11 (Mical-i + SelR-i) embryos were used for quantification. Error bars represent ± SEM. Kruskal-Wallis tests were performed in K–N, R, and V: *P < 0.05, **P < 0.01, ***P < 0.001, ****P < 0.0001, and ns is not significant.

### Blocking Methionine 44 oxidation exhibits similar phenotypes to those of Mical knockdown

Previous studies showed that redox reactions at the Methionine 44 residue (M44) of the major Drosophila cytoplasmic actin (Actin5C), mediated by Mical/SelR, regulate F-actin polymerization and depolymerization (*27, 31*). We generated wildtype and non-oxidizable versions of actin: Act5C-WT and Act5C-M44L, respectively; see Methods). We used these lines to determine how actin modifications on M44 affect actin dynamics and cell wound repair by generating the wounds in embryos overexpressing wild-type or M44L-actin with an F-actin reporter in a heterozygous Act5C deletion (Act5C^Δ^) background. Upon laser wounding, a robust actin ring formed in embryos expressing Act5C-WT, whereas expression of Act5C-M44L led to phenotypes similar to those observed in Mical knockdown embryos: premature actin ring disassembly, actin accumulation inside the wounds, slower wound contraction, and decreased actin ring width/intensity (Fig. 5, A-G; Table S2). Thus, actin modifications at M44 are required for proper F-actin dynamics, the formation of a robust actin ring at the wound periphery, and efficient wound repair.

**Figure 5.**
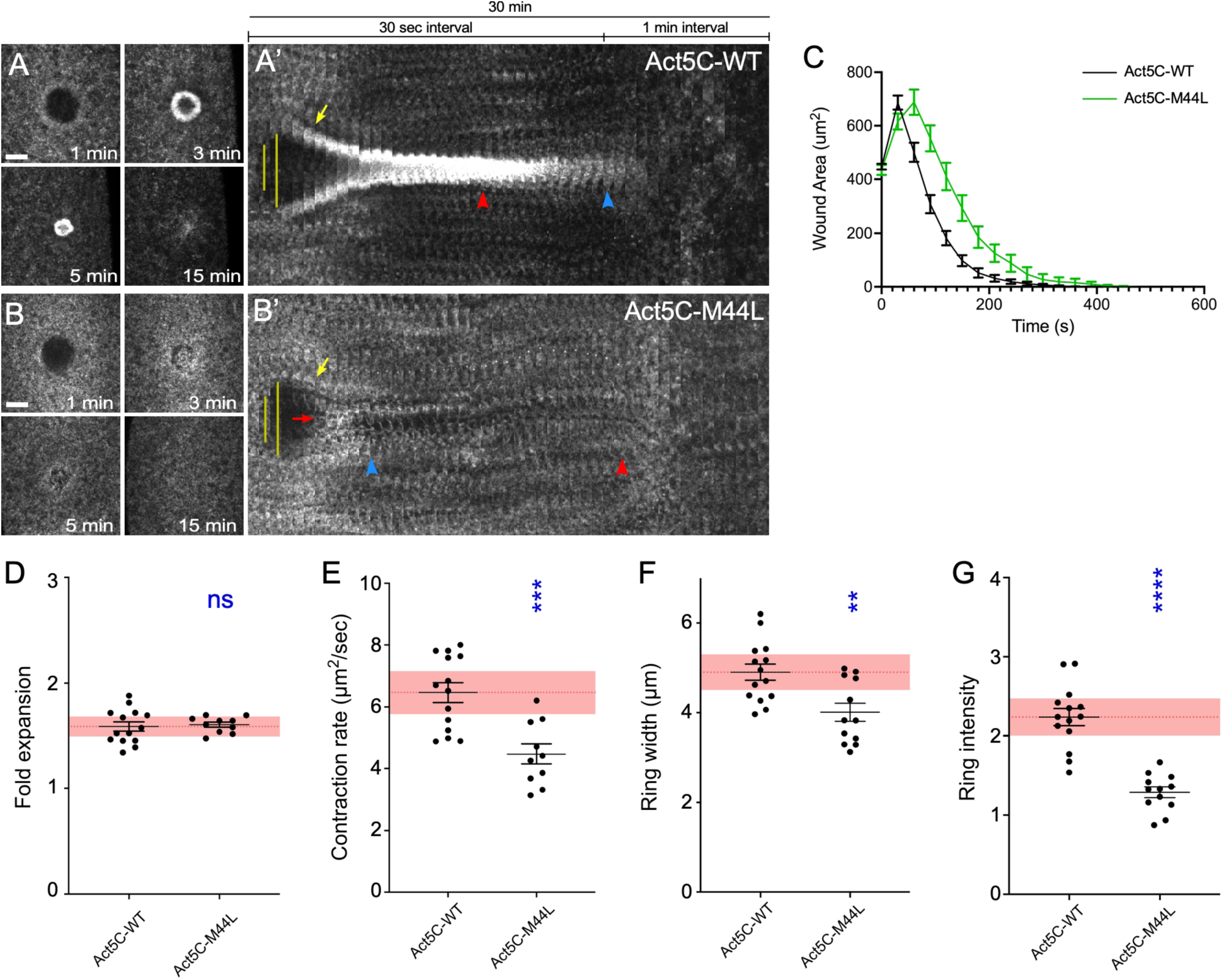
Blocking oxidation at Methionine 44 of actin disrupts actin dynamics during cell wound repair. (A-B) Actin dynamics (sGMCA) during cell wound repair in NC4–6 staged Drosophila embryos: control (wild type Act5C) (A), and Act5C-M44L (2) (B). (A’-B) Kymographs across the wound area depicted in A-BE, respectively. Wound expansion is highlighted by yellow lines. Actin recruitment to the actomyosin ring is indicated by yellow arrows. Actin accumulation internal to the wound is indicated by red arrows. Wound closure is indicated by red arrowheads. Actomyosin ring disassembly (or lack thereof) is indicated by blue arrowheads. Scale bar: 20 µm. Time after wounding is indicated. (C) Quantification of the wound area over time for control (n = 14) and Act5C-M44L (n = 10). (D–G) Quantification of fold wound expansion (D), contraction rate (E), actin ring width (F), and actin ring intensity (G) for control and Act5C-M44L. Error bars represent ± SEM. Mann-Whitney tests were performed in D–G: ****P < 0.0001.: *P < 0.05, **P < 0.01, ***P < 0.001, ****P < 0.0001, ns is not significant.

### F-actin architectures in Act5C-M44L are not identical to those in Mical knockdowns

If Mical is the only protein oxidizing Actin5C, we expected that F-actin architectures in embryos expressing Act5C-M44L would have the same features as those in Mical knockdown embryos. Using super-resolution microscopy, we generated wounds in embryos expressing Act5C-WT and Act5C-M44L with an F-actin reporter. All Haralick features in embryos expressing Act5C-M44L are similar to those in Mical knockdown embryos: lower VARIANCE and higher CORRELATION, UNIFORMITY, and HOMOGENEITY (Fig. 4, K-N and 6, A-J; Table S3). Surprisingly, while Mical knockdown embryos exhibited a significant increase of F-actin with DV axis orientation in the actin ring (Fig. 4, O-V), embryos expressing Act5C-M44L exhibit a significant increase of F-actin with AP axis orientation (Fig. 6, K-N).

**Figure 6.**
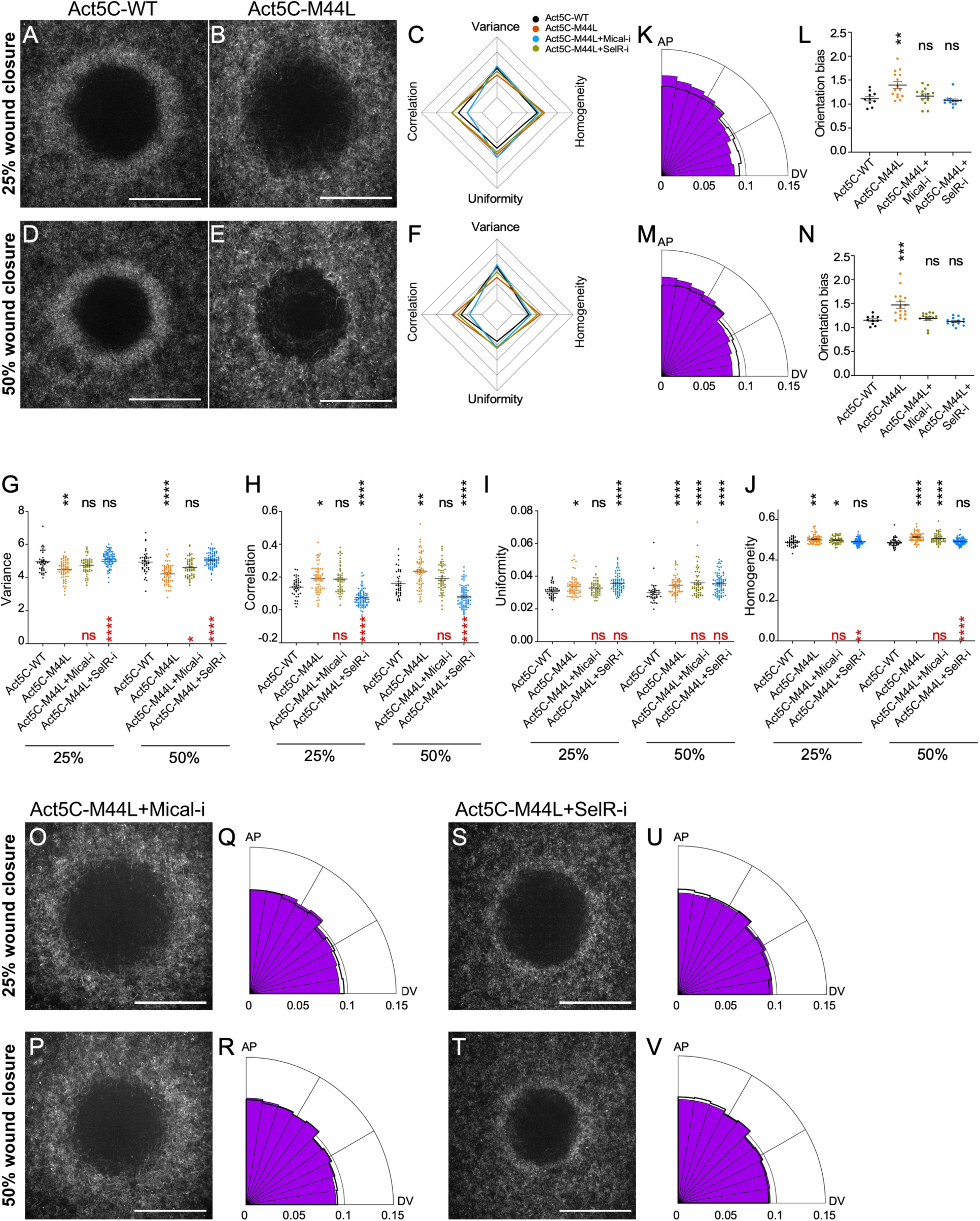
Blocking oxidation at Methionine 44 of actin does not mimic Mical knockdown phenotypes. (A-B) Deconvolved super-resolution confocal images from NC4–6 Drosophila embryos expressing sGMCA at the time point corresponding to 25% of max wound area: control (Act5C-WT) (A) and Act5C-M44L (B). (C) Radar chart of average Haralick features comparing control, Act5C-M44L, Act5C-M44L + Mical-i, and Act5C-M44L + SelR-i at the time point corresponding to 25% of the max wound area. (G-J) Quantification of Haralick features comparing control, Act5C-M44L, Act5C-M44L + Mical-i, and Act5C-M44L + SelR-i at time points corresponding to 25% and 50% of max wound area. (D-E) Deconvolved super-resolution confocal images from NC4–6 Drosophila embryos expressing sGMCA at the time point corresponding to 50% of max wound area: control (Act5C-WT) (D) and Act5C-M44L (E). (F) Radar chart of average Haralick features comparing control, Act5C-M44L, Act5C-M44L + Mical-i, and Act5C-M44L + SelR-i at the time point corresponding to 50% of the max wound area. (K) Radial histograms showing the distribution of actin filament orientations ranging from anterior–posterior (AP) bias to dorsal–ventral (DV) bias in Act5C-M44L at the time point corresponding to 25% of max wound area. The black line indicates the control distribution. (L) Quantification of the distribution of actin filament orientations comparing control, Act5C-M44L, Act5C-M44L + Mical-i, and Act5C-M44L + SelR-i at the time point corresponding to 25% of max wound area. (M) Radial histograms showing the distribution of actin filament orientations ranging from anterior–posterior (AP) bias to dorsal–ventral (DV) bias in Act5C-M44L at the time point corresponding to 50% of max wound area. The black line indicates the control distribution. (N) Quantification of the distribution of actin filament orientations comparing control, Act5C-M44L, Act5C-M44L + Mical-i, and Act5C-M44L + SelR-i at the time point corresponding to 50% of max wound area. (O-P) Super-resolution confocal images from NC4–6 Drosophila embryos expressing sGMCA at time points corresponding to 25% (O) and 50% (P) of max wound area: Act5C-M44L + Mical-i. (Q-R) Radial histograms showing the distribution of actin filament orientations ranging from anterior–posterior (AP) bias to dorsal–ventral (DV) bias in Act5C-M44L + Mical-i at time points corresponding to 25% (Q) and 50% (R) of max wound area. The black line indicates the control distribution. (S-T) Super-resolution confocal images from NC4–6 Drosophila embryos expressing sGMCA at time points corresponding to 25% (S) and 50% (T) of max wound area: Act5C-M44L + SelR-i. (U-V) Radial histograms showing the distribution of actin filament orientations ranging from anterior–posterior (AP) bias to dorsal–ventral (DV) bias in Act5C-M44L + SelR-i at time points corresponding to 25% (U) and 50% (V) of max wound area. The black line indicates the control distribution. Scale bar: 20 µm. 10 (Act5C-WT), 16 (Act5C-M44L), 14 (Act5C-M44L + Mical-i), and 120 (Act5C-M44L + SelR-i) embryos were used for quantification. Error bars represent ± SEM. Kruskal-Wallis tests were performed in G-H, L, and N: *P < 0.05, **P < 0.01, ***P < 0.001, ****P < 0.0001, and ns is not significant.

**Figure 7.**
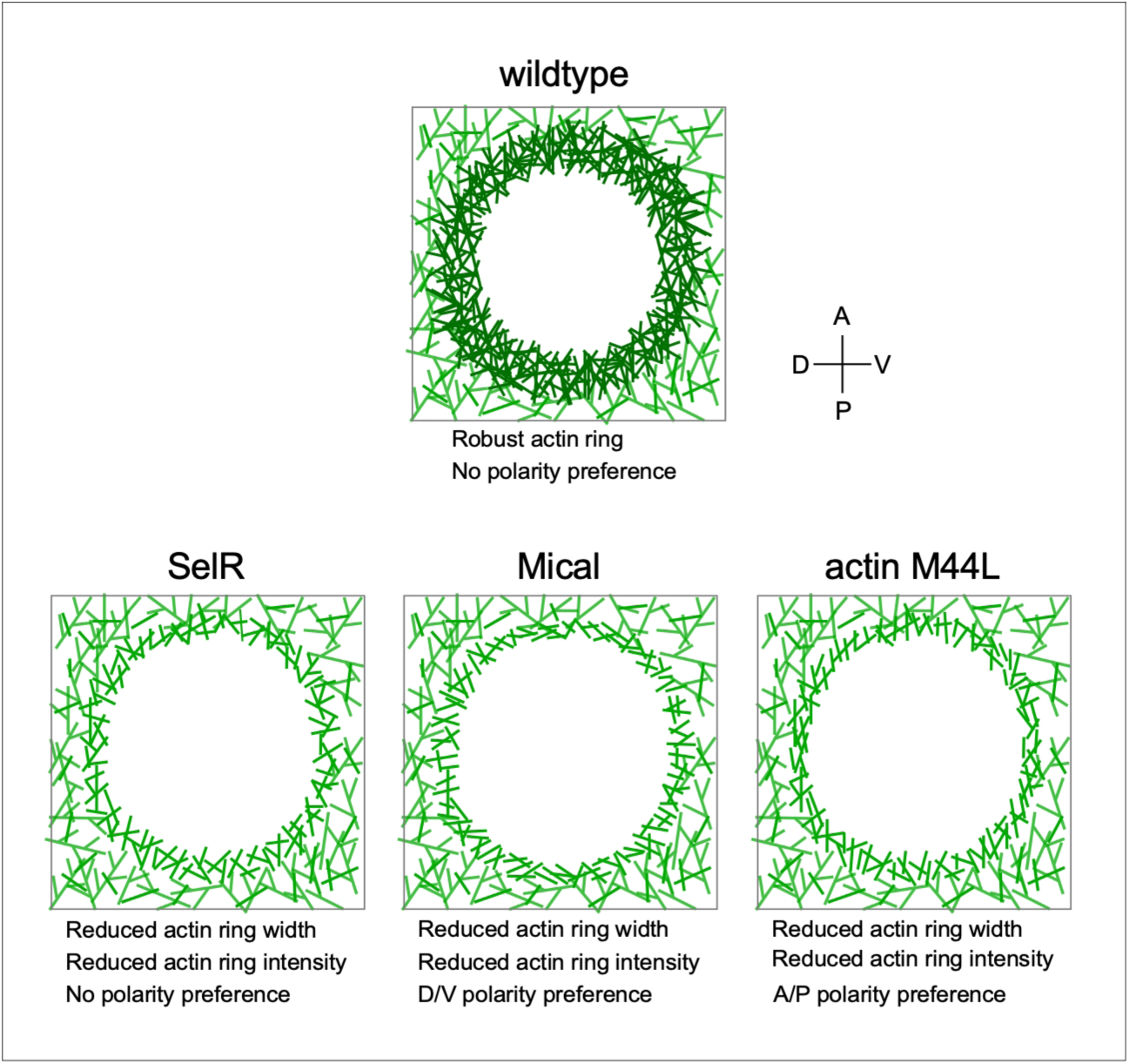
Actin redox functions affecting actin ring dynamics during cell wound repair. Schematic diagram summarizing actin ring architectures in wild-type, SelR and Mical knockdowns, and the actin M44L mutation. In wild-type, the actin ring is highly dense and is formed by a uniform distribution of F-actin orientations. SelR knockdowns, where actin reduction was decreased, exhibit reduced actin ring width and intensity, and no preference for F-actin orientation. Mical knockdowns, in which actin oxidation was decreased, exhibit reduced actin ring width and intensity, and actin filaments within the actin ring are oriented along the D/V axis. Actin M44L, which blocks the Mical-dependent actin oxidation site, exhibits reduced actin ring width and intensity, and actin filaments within the actin ring are oriented along the A/P axis.

Several possibilities could account for the opposite F-actin orientations observed: 1) Mical oxidases other proteins in addition to actin during wound repair, 2) M44 in actin is oxidized by more than one enzyme, and 3) both 1) and 2). To test these possibilities, we knocked down Mical in embryos expressing Act5C-M44L and generated wounds on which we performed texture and F-actin orientation analyses in the actin ring. If Mical oxidases additional proteins to actin, we would expect to observe phenotypes similar to those of the Mical knockdown. If actin M44 is oxidized by more than one enzyme, we would expect to observe phenotypes similar to those of Act5C-M44L alone. If the phenotypes are not similar to either Mical knockdown or Act5C-M44L, both possibilities are occurring in the context of cell wound repair. While Mical knockdown and Act5C-M44L alone both exhibit similar Haralick features (Fig. 4, K-N and 6, G-J), when combined they exhibit no significant differences in VARIANCE and CORRELATION (Fig 6, G, H, O, and P; Table S3). In addition, F-actin orientations in Mical knockdown with Act5C-M44L exhibit a similar bias to those of control (Fig. 6, L, N, Q, and R; Table S3).

Since double knockdown of Mical and SelR partially rescues their individual phenotypes (Fig. 3 and 4), we examined whether knocking down SelR in the Act5C-M44L background can similarly rescue the phenotypes of Act5C-M44L. SelR knockdown in the Act5C-M44L background results in a more dense actin ring formation compared with Act5C-M44L as previously observed in Mical and SelR double knockdown and partially rescues the phenotypes (Fig. 4, and 6, B, E-J, S-V, L, and N; Table S3). Intriguingly, when comparing Haralick features between the SelR knockdown with Act5C-M44L and the Mical+SelR double knockdown, the SelR knockdown with Act5C-M44L shows lower CORRELATION values (Fig. 4, L and 5, H). Thus, our results suggest that oxidation of actin M44 is regulated by more than one oxidation enzyme, and that, in addition to actin, Mical and SelR regulate redox modifications of other proteins during cell wound repair.

## DISCUSSION

Cell wound repair is a highly conserved process that enables all organisms to protect themselves against daily damage caused by physiological and environmental stresses. Actomyosin ring contraction at the wound periphery is a major mechanism that physically closes cell wounds in many cell wound repair models. Here, we show that reversible actin modifications mediated by the Rab35/Mical/SelR pathway regulate actin ring assembly and disassembly during cell wound repair. While the wound phenotypes of Mical and SelR share many similarities, they differ in their effects on F-actin architecture at the wound edge. Interestingly, our findings suggest that Mical oxidizes more than actin, and that actin is oxidized by more enzymes than Mical.

### Rab35 regulates actin redox through Mical and SelR for actin ring assembly and disassembly during cell wound repair

While the role of Rab GTPases in regulating membrane trafficking is well known, Rab35 also regulates the actin cytoskeleton in cellular processes, such as cytokinesis. Cytokinesis requires the formation, contraction, and remodeling of a robust actin ring, which is also required for efficient cell wound repair. During cytokinesis, Rab35 recruits Mical to the intercellular bridge, where Mical-mediated actin oxidation leads to actomyosin ring disassembly and successful abscission (*29*) SelR reverses these modifications (*31, 32, 37*). Hence, Mical depletion slows actomyosin ring clearance in the intercellular bridge, while SelR depletion accelerates it (*29, 32*). We find that both Mical and SelR recruitment to cell wounds is regulated by Rab35, suggesting that Rab35 is a key regulator of dynamic actin redox modifications during the repair process. In contrast to cytokinesis, we find that Mical and SelR knockdowns dynamically affect both the formation and remodeling of the actomyosin ring during cell wound repair. In particular, we find that Mical depletion resulted in a pronounced D/V polarity preference for actin filaments at the wound edge, whereas SelR results in a uniform distribution of actin filament polarities. Interestingly, double knockdown of Mical + SelR rescued the D/V polarity difference of Mical resulting in a uniform distribution of actin filament polarity at the wound edge. Thus, proper actomyosin ring assembly and disassembly require a dynamic balance between actin oxidation and reduction, rather than the activity of either enzyme alone. In addition, Mical-mediated actin oxidation has been shown to cooperate with actin-binding proteins such as Profilin and cofilin to enhance filament turnover and remodeling (*38–40*). As such, Rab35-dependent recruitment of Mical and SelR during cell wound repair could influence multiple layers of actin regulation. Further studies are needed to investigate how Rab35 dynamically regulates the activity balance between Mical and SelR across the different steps of the repair process.

### Actin oxidation regulates proper F-actin orientations to form a robust actomyosin ring

Mical-mediated oxidation leads to F-actin disassembly, whereas SelR restores F-actin polymerization by reducing oxidized actin. Hence, reversible actin redox by Mical and SelR is thought to control F-actin length and turnover (*41–43*). Our super-resolution microscopy analyses indicates that actin oxidation also contributes to proper F-actin orientation within the actomyosin ring. Interestingly, while Mical knockdowns exhibit a D/V polarity preference for actin filaments at the wound edge, the actin M44L point mutation exhibits the opposite A/P preference for actin filament orientation. A combination of Mical knockdown and the actin M44L mutation returns the actin filament orientation to a uniform distribution of polarities. These results suggest that Mical likely oxidizes more proteins than actin, and that actin is likely oxidized by more proteins than Mical. Consistent with this, while Actin M44 in known to be a major functional target of Mical, actin has been shown to be oxidized on other amino acid residues (*44*). In addition, a previous study showed that MICAL-2 oxidizes Arp3B to enhance branched actin network disassembly (*45*). Intriguingly, the A/P actin filament orientation preference of the Actin M44L mutation is similar to that we have shown previously for Arp3 knockdowns (*36*). This phenotype similarity suggests that Actin M44 may be involved in branched actin network formation. Indeed, previous Cryo-EM studies have shown that the actin D-loop region, including Actin M44, is a docking site for Arp3 to polymerize daughter filaments (*46–49*). Actin M44 oxidation could affect the polymerization of daughter branched filaments, which in turn, would influence F-actin orientation. The regulation of branched F-actin networks is an alternative possibility for why Actin M44L exhibits the opposite F-actin orientation to Mical knockdown, since Mical could oxidize Arp3 as well as actin. Overall, our results suggest that reversible actin redox regulates more than F-actin assembly and disassembly to form a robust actomyosin ring during cell wound repair.

### Reversible actin modifications regulate dynamic actin architectures

Posttranslational modifications (PTMs) are central drivers of dynamic protein functions across diverse cellular processes (*50–53*). Just as histone and tubulin PTMs have been extensively characterized as regulatory “codes”, actin PTMs have similarly emerged as fundamental regulators of actin’s roles in cell behaviors and development (*42, 44, 54–57*). Notably, N-terminal acetylation of actin has been shown to be essential for filopodia and lamellipodia formation during cell migration (*58*). Although multiple actin modification sites have been identified previously, their spatial and temporal regulation—and their full physiological significance—remain incompletely understood. Here, we showed that reversible actin redox modifications regulate multiple aspects of actomyosin ring functions during cell wound repair: F-actin architecture that affects actomyosin ring assembly and contraction necessary for efficient wound closure, and remodeling of the actomyosin ring to restore the cell cortex to its unwounded state. Actin architectures are highly complex and change dynamically across biological processes, from cells to organs, including cell migration, wound repair, and normal morphogenesis. Actin modifications provide an elegant means in which to regulate complex F-actin networks that carry out dynamic spatial and temporal orchestrations during these events.

## ACKNOWLEDGEMENTS

We thank Viktor Stjepić, Tony Cooke, the Bloomington Stock Center, the Kyoto Stock Center, the Harvard Transgenic RNAi Project, FlyBase, the Fred Hutch/Leica Center of Excellence, and the Drosophila Genome Resource Center for advice, microscopes, DNAs, flies, and other reagents used in this study.

This work was supported by NIH grant R35GM161275 (to SMP), the Mark Groudine Chair for Outstanding Achievements in Science and Service (to SMP), and the NCI Cancer Center Support Grant P30 CA015704 (for Shared Resources). The funders had no role in study design, data collection and interpretation, or the decision to submit the work for publication.

The authors declare no competing financial interests.

All data needed to evaluate the conclusions in the paper are present in the paper and/or the Supplementary Materials.

## Author Contributions

All authors contributed to the design of the experiments, performed experiments, analyzed data, and performed the morphometric analyses. MN wrote the manuscript with input from all authors.

## MATERIALS AND METHODS

Reagents used in this study are described in Table S1.

### Fly stocks and genetics

Flies were cultured and crossed at 25°C on yeast-cornmeal-molasses-malt medium. Flies used in this study are described in Table S1. All fly stocks were treated with tetracycline, then tested by PCR to ensure that they did not harbor Wolbachia.

An actin reporter, sGMCA (spaghetti squash driven, moesin-alpha-helical-coiled and actin binding site fused to GFP) reporter (*59*) or the mScarlet-i (sStMCA; (*60*)) fluorescent equivalent was used to follow wound repair dynamics of the cortical cytoskeleton.

To knockdown genes, RNAi lines were driven maternally using the GAL4-UAS system with P{matalpha4-GAL-VP16}V37 (Bloomington #7063 (*35*)) for driving RNAi or P{w[+mC]=GAL4::VP16-nos.UTR}MVD1 (Bloomington #4937) for driving Actin5C and Actin5C-M44L in heterozygous Act5C mutant backgrounds. Localization patterns and mutant analyses were performed at least twice from independent genetic crosses and ≥10 embryos were examined unless otherwise noted. Images representing the average phenotype were selected for figures.

### Generation of mScarlet-SelR reporter

To generate sqh-StFP (mScarlet-i) -SelR, the SelR ORF was amplified from BDGP clone RE73235 and fused 5′ to StFP. The resulting StFP-SelR fusion was cloned into pSqh5′+3′UTR (*34*) as a 5′ StuI-3′ XbaI fragment. To generate transgenic flies, sqh-StFP-SelR construct (500 µg/ml) was injected along with the pTURBO helper plasmid (100 µg/ml) into isogenic w* flies as previously described (*61*).

### Generation of Mical and SelR RNAi lines

RNAi lines for Mical and SelR were generated using the method previously described (*62, 63*). 2 oligos were annealed and cloned into pUASzMIR (DGRC_1432). UASz-Mical RNAi(2) and UASz-SelR RNAi(1) were injected into M{3xP3-RFP.attP}ZH-86Fb (BDSC_24749).

### Generation of Actin5C mutant

Act5C mutant (Act5C^Δ^) was generated using the CRISPR-Cas9 system. Two gRNAs (5’-gagaatttgcgtggtttccttgg-3’ and 5’- ctgggcaagaggatcaggatcgg-3’) were cloned into pCFD5 vector ((*64*); Addgene#73914) as two tRNA-flanked gRNAs and then the resulting vector was injected into the Cas9 fly embryos (BDSC #78782 that had been crossed into a w- background). The deletion of the entire Act5C CDS was confirmed by PCR and sequencing.

### Generation of the Actin5C point mutation

To generate a point mutation on Act5C M44 as previously described (*27*), we used two primers (5’-tcaccagggtgtgttggtcggcatggg-3’ and 5’-cccatgccgaccaacacaccctggtga-3’). Full-length Act5C-WT and Act5C-M44L were amplified by PCR and then cloned into the 3xUASz-p10 vector (*36*). 3xUASz-Act5C-WT-p10 and 3xUASz-Act5C-M44L (500 µg/ml) were injected along with the pTURBO helper plasmid (100 µg/ml) into isogenic w* flies as previously described (*61*).

### Embryo handling and preparation

Nuclear cycle (NC) 4-6 *Drosophila* embryos were collected from 0-30 min at room temperature (22°C). Embryos were hand dechorionated, placed onto No. 1.5 coverslips coated with glue, and covered with Series 700 halocarbon oil (Halocarbon Products Corp).

### Laser wounding

All wounds were generated with a pulsed nitrogen N2 Micropoint laser (Andor Technology Ltd., Concord, MA, USA) tuned to 435 nm and focused on the cortical surface of the embryo. A region of interest was selected in the lateral midsection of the embryo and ablation was controlled by MetaMorph. On average, ablation time was less than 3s, and time-lapse imaging was initiated immediately. Occasionally, a faint grid pattern of fluorescent dots is visible at the center of wounds that arises from damage to the vitelline membrane that covers embryos.

### Microscopy

All imaging was performed at room temperature (22°C). For live imaging, the following microscope was used: Revolution WD systems (Andor Technology Ltd., Concord, MA, USA) mounted on a Leica DMi8 (Leica Microsystems Inc., Buffalo Grove, IL, USA) with a 63x/1.4 NA objective lens and controlled by MetaMorph software. Images and videos were acquired with 488 nm and 561 nm, using an Andor iXon Ultra 897 or 888 EMCCD cameras (Andor Technology Ltd., Concord, MA, USA). All images for cell wound repair were 17-20 µm stacks/0.25 µm steps. For single color, images were acquired every 30 sec for 15 min and then every 60 sec for 25 min. For dual green and red colors, images were acquired every 30 sec for 30 min. Due to rapid bleaching, YFP-Rab35 and sStMCA images were acquired every 1 min for 5 min and then every 3 min for 39 min.

### Live super-resolution microscopy

Super-resolution imaging was performed using a VT-iSIM (VisiTech International) mounted on a Leica DMi8 (Leica Microsystems) with a 100×/1.4 NA objective lens under the control of MetaMorph software (Molecular Devices). Images were acquired using a 488-nm laser using an Orca-Fusion C14440-20UP camera (Hamamatsu Photonics). Time-lapse images were acquired with 10-µm stacks/0.25-µm steps every 15 s for 7.5 min followed by every 1 min for 25 min and deconvolved using Huygens software (Scientific Volume Imaging).

### Image processing, analysis, and quantification

All images were analyzed with Fiji (*65*). Measurements of wound area were done manually. To generate xy kymographs, all time-lapse xy images were cropped to 5.8 µm x 94.9 µm and then each cropped image was lined up. For fluorescent line plots, the mean fluorescence profile intensities were calculated from 51 equally spaced radial profiles anchored at the center of the wound, swept from 0° to 180°. Radial profiles of diameter 301 pixels were used. Fluorescence intensity profiles were calculated and averaged using an in-house code ((*36*): available at https://github.com/FredHutch/wound_radial_lineplot) using MATLAB R2020b (MathWorks). For dynamic line plots, we generated fluorescent profile plots from each time point and then concatenated them. The lines represent the averaged fluorescent intensity and the gray area is the 95% confidence interval.

Quantification of the width and average intensity of actin ring, wound expansion, and closure rate was performed as follows: the width of actin ring was calculated with two measurement, the ferret diameters of the outer and inner edge of actin ring at 90 sec post-wounding. Using these measurements, the width of actin ring was calculated with (outer ferret diameter – inner ferret dimeter)/2. The average intensity of actin ring was calculated with two measurement. Instead of measuring ferret diameters, we measured area and integrated intensity in same regions as described in ring width. Using these measurements, the average intensity in the actin ring was calculated with (outer integrated intensity - inner integrated intensity)/(outer area - inner area). To calculate relative intensity for unwounded (UW) time point, average intensity at UW was measured with 50x50 pixels at the center of embryos and then averaged intensity of actin ring at each timepoint was divided by average intensity of UW. Wound expansion was calculated with max wound area/initial wound size. Closure rate was calculated with two time points, one is t_max_ that is the time of reaching maximum wound area, the other is t<half that is the time of reaching 50-35% size of max wound since the slope of wound area curve changes after t<half. Using these time points, average speed was calculated with (wound area at t_max_ – wound area at t<half)/t_max_-t<half.

To quantify Mical and SelR recruitment to wounds, we drew a 5-pixel-wide line along the wound edge at the 2-min post-wounding time point and measured the average intensity. Using the same region at the 2 min post-wounding, the average intensity at the unwounded time point was measured. Then the average intensity at the 2-minute post-wounding time point was divided by that at the unwounded time point.

Texture analysis was assessed using cropped sections of deconvolved single z-slice images 0.25–0.5 μm from the surface of the cortex of the actin ring at 70% wound closure. The gray-level cooccurrence matrix of eight 50 × 50–pixel cropped areas of the wound edge was used to compute four texture-related statistical properties per embryo, variance/contrast (the intensity between a pixel and its neighbor), correlation (linear dependency), uniformity (constancy of intensity level distribution), and homogeneity (the tightness of distribution; (*66*)), similar to those previously described (*60*). Textural features were extracted using MATLAB R2020b (MathWorks) and were graphed on radar charts and dot plots using R.

Filament orientation was determined using a method similar to that described previously (*36, 67*) using Matlab 2020b (MathWorks). The images were convolved using the Sobel operator and the gradient intensity and gradient direction from the DV axis were retrieved. The orthogonal direction of the intensity gradient is the filament orientation. To focus on the filaments at the edge of the wound, we limited the analysis to the pixels more than 150 pixels and less than 200 pixels from the center of the wound. Additionally, we placed a threshold based on the magnitude of the intensity gradient to remove background noise. The actin filament distribution was plotted on a radial histogram for visualization. Additionally, the orientation bias was calculated as a ratio of the density of filaments with orientations between 80°–90° and 0°–10°. This was calculated and graphed in R.

### qPCR

Total RNA was obtained from 100 embryos (0-30 min old) using TRIzol (Invitrogen). 1 µg of total RNA was used for reverse transcription with the iScript™ gDNA Clear cDNA Synthesis Kit (Bio-Rad). RT-PCR analysis was performed using the iTaq™ Universal SYBR® Green Supermix (Bio-Rad) with two individual parent sets and two technical replicates on the CFX96TM Real Time PCR Detection System (Bio-Rad). RpL32 was used as a reference gene. The % knockdown was calculated using the ΔΔCq calculation method compared with control (vermilion knockdown). Same primer sets RpL32 was used in previously described (*36*).

### Statistical analysis

All statistical analyses were done using Prism 8 (GraphPad, San Diego, CA). Gene knockdowns were compared to the appropriate control, and statistical significance was calculated using a Mann-Whitney test or Kruskal-Wallis test with *P*<0.05 considered significant.

